# Mutational meltdown of microbial altruists in *Streptomyces coelicolor* colonies

**DOI:** 10.1101/2020.10.20.347344

**Authors:** Zheren Zhang, Shraddha Shitut, Bart Claushuis, Dennis Claessen, Daniel E. Rozen

## Abstract

In colonies of the filamentous multicellular bacterium *Streptomyces coelicolor*, a sub-population of cells arise that hyper-produce metabolically costly antibiotics, resulting in division of labor that maximizes colony fitness. Because these cells contain large genomic deletions that cause massive reductions to individual fitness, their behavior is altruistic, much like worker castes in social insects or somatic cells in multicellular organisms. To understand the reproductive and genomic fate of these mutant cells after their emergence, we use experimental evolution by serially transferring populations via spore-to-spore transfer for 25 cycles, reflective of the natural mode of bottlenecked transmission for these spore-forming bacteria. We show that, in contrast to wild-type cells, altruistic mutant cells continue to significantly decline in fitness during transfer while they delete larger and larger fragments from their chromosome ends. In addition, altruistic mutants acquire a roughly 10-fold increase in their base-substitution rates possibly due to mutations in genes for DNA replication and repair. Ecological damage, caused by reduced sporulation, coupled with irreversible DNA damage due to point mutations and deletions, leads to an inevitable and irreversible type of mutational meltdown in these cells. Taken together, these results suggest that the altruistic cells arising in this division of labor are analogous to reproductively sterile castes of social insects.

## Introduction

Multicellular organisms show enormous variation in size and complexity, ranging from multicellular microbes to sequoias and whales, and from transient undifferentiated cellular clusters to stable individuals with highly specialized cell types. Despite their differences, a recent study showed that a central factor determining organismal complexity is the way in which multicellular organisms are formed ^1^. Clonal groups, where relatedness among cells is high, show more cellular specialization and an increased likelihood of expressing a reproductive division of labor between somatic and germ cells ^1–4^. By contrast, groups with aggregative multicellularity like dictyostelid social amoebae or myxobacteria, which potentially have lower relatedness between cells if unrelated genotypes coaggregate during development, tend to show reduced specialization ^5–7^. Thus, in analogy with sterile castes of workers within colonies of social insects that are morphologically differentiated to perform specialized tasks, the extreme altruism needed for reproductive sterility is facilitated by high relatedness^8^.

In microbes, the requirement of high relatedness is most easily met if colonies are initiated from a single cell or spore. High relatedness during multicellular growth or development is even further guaranteed if the cells within colonies remain physically connected to each other, as observed in filamentous streptomycetes ^9,10^. These bacteria have a well-characterized reproductive division of labour due to a developmental program that leads to the formation of durable spores following a period of vegetative growth and the elaboration of spore-bearing aerial hyphae ^11,12^. In addition, we recently showed that colonies are further divided into a sub-population of cells that hyper-produces antibiotics ^13^. The ability to generate this sub-population of cells benefits the whole colony, by virtue of their increased antibiotic production, without causing a decline in overall colony spore production. However, the fitness of the mutant sub-population is significantly reduced. Here we provide a detailed examination of the evolutionary fate of these specialized cells and provide evidence that they represent a terminally differentiated altruistic cell type within these multicellular microbes.

*Streptomyces* are bacteria that live in the soil and produce a broad diversity of antibacterial and antifungal compounds, among other specialized metabolites ^14,15^. Division of labor allows *Streptomyces coelicolor* colonies to partly offset the metabolic cost of producing these compounds. However, differentiation into this hyper-producing cell type is accompanied by huge fitness costs due to massive deletions of up to 1 Mb from the ends of their linear chromosomes. Examining independent mutant strains, we found a strong positive correlation between the size of genome deletions and the amount of antibiotics produced, as well as a strong negative correlation between deletion size and spore production. In addition, as shown earlier^13^ and in Fig. 1A, competition assays revealed that mutant strains were strongly disadvantaged. Indeed, even when the initial frequency of mutants in mixed colonies was as high as ~80%, their final frequency declined to less than 1% after one cycle of colony growth. These results indicate that mutant strains would be quickly eliminated during competitive growth. We hypothesized that, like sterile insect workers or somatic cells that are differentiated to perform specialized tasks, these altruistic cells represent a sterile microbial caste because they are functionally cohesive, showing uniformly increased antibiotic production, and arise due to a shared mechanism of genome deletions. However, as our results were based on static colonies, we lacked insight into the fate of these cells after they emerged.

**Fig. 1.**
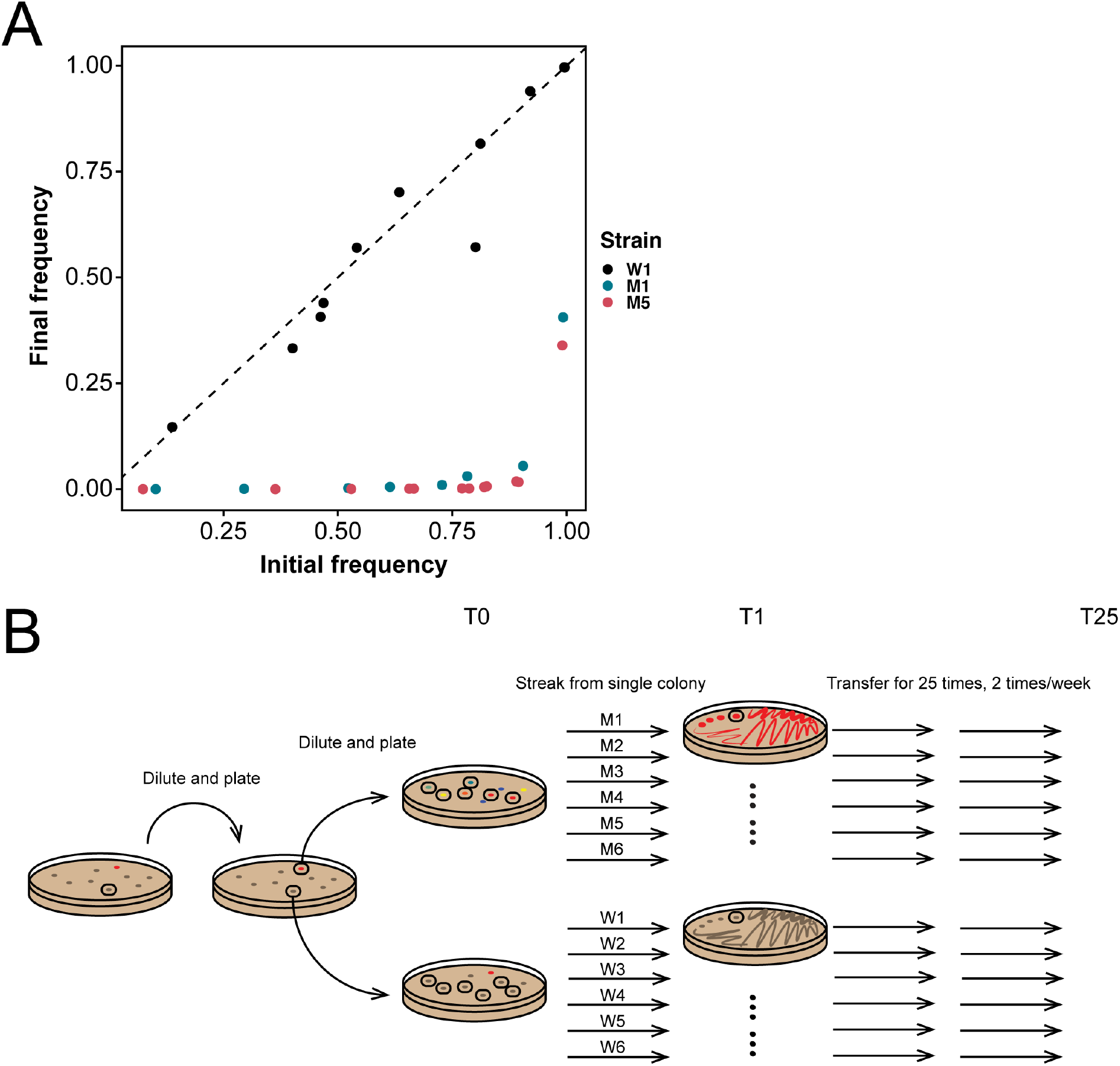
Overview of the experimental design. (**A**) Initial and final frequency of three T0 strains from different lineages during competition with the WT ancestor. The dashed line shows the expectation if initial and final frequencies are equal, as seen for the strain from the WT lineage (W1). By contrast, mutant fitness (M1 and M5) is dramatically lower than the WT, dropping to < 1% even when starting from as high as approximately 73% (M1) or 82% (M5). (**B**) The schematic of our experimental setup. An ancestral WT colony was picked and plated to obtain individual colonies. One mutant and one WT colony were picked and plated to obtain six WT and six mutants clones. Lineages were subsequently transferred via single colony bottlenecks for 25 transfers. Each timepoint of these transfers is designated as T plus the transfer number.

To address this question, the current study tracked the fate and fitness of altruistic mutant and wildtype lineages during short-term experimental evolution. To reflect the manner of spore-to-spore reproduction in these bacteria, lineages were serially transferred via single colonies using a mutation accumulation design ^16^ (Fig. 1B). In contrast to much longer-term experiments using this approach in other microbes, where fitness declines extremely slowly ^17,18^, we observed significant fitness reductions, including extinction, in our mutant lineages after only 25 transfers. These changes were not only associated with continued deletions to the chromosome ends, but also the tendency for lineages to become hypermutators likely due to errors in genes for DNA replication and repair ^19,20^. Together these data support the idea that this specialized sub-population of cells within *Streptomyces* colonies are analogous to a functionally differentiated sterile caste that persist due to their inclusive fitness benefits to the entire colony and further highlights the idea that clonal propagation and high relatedness can give rise to a broad diversity of functionally specialized cells within these multicellular bacteria.

## Results

### Phenotypic changes during serial transfer

To track the fate of different mutant lineages harboring different spontaneous genomic deletions we transferred six WT (W1-W6) and six mutant (M1-M6) strains for 25 transfers using a mutation accumulation design^16^ through single spore bottlenecks twice per week (Fig 1B). Details of strain origins are given in the methods. Consistent with our earlier results ^13^, we first confirmed that the starting competitive fitness of a subset of these mutants was significantly reduced compared to the WT ancestor (Fig 1A). Even when mutant lineages were inoculated at an initial frequency as high as roughly 80%, their final frequency during paired competition declined to less than 1%. In addition, the mutant strains that were used to initiate the MA experiment produced significantly fewer colony-forming unit (CFU) after clonal development than their WT counterparts (Wilcoxon rank sum test, *P* = 0.0022, Fig. 3A). Strains were sampled every approximately 5 transfers, with the exception of one WT lineage (W3) that was sampled more frequently after it acquired chromosome deletions after the 7^th^ transfer (as described below). One of the six mutant lineages (M2) acquired a bald morphology after the 5^th^ transfer and became functionally extinct due to a total loss of spore production and was not included in fitness analyses (Fig. S1).

To identify phenotypic changes in evolved lineages, we screened for two easily scored traits that are indicative of deletions to the right chromosome arm ^13^. Chloramphenicol susceptibility, due to the deletion of *cmlR1* (SCO7526)/*cmlR2* (SCO7662), indicates a deletion of at least 322 kb ^21,22^ and arginine auxotrophy, due to the deletion of *argG* (SCO7036), corresponds to a deletion of at least 843 kb ^23^. In addition, we analyzed changes to resistance to three other antibiotics. As is evident in Fig. 2A, whereas the WT lineages remained resistant to chloramphenicol (except for W3) the minimal inhibitory concentration (MIC) of mutant lineages were lower than the WT or declined during the course of the experiment. On the basis of these results, W3 was hereafter analyzed as a mutant lineage, despite its WT origin. A trend towards increased arginine auxotrophy was also observed in mutant lineages (Fig. 2B), suggesting that continuous chromosome deletions occurred during the course of the experiment (genomic changes are described below). Tests for susceptibility to other antibiotics (Fig. S2) also showed similar trends as those found for chloramphenicol, with the exception of the bald populations from M2 that showed a 4-fold increase in the MIC for ciprofloxacin.

**Fig. 2.**
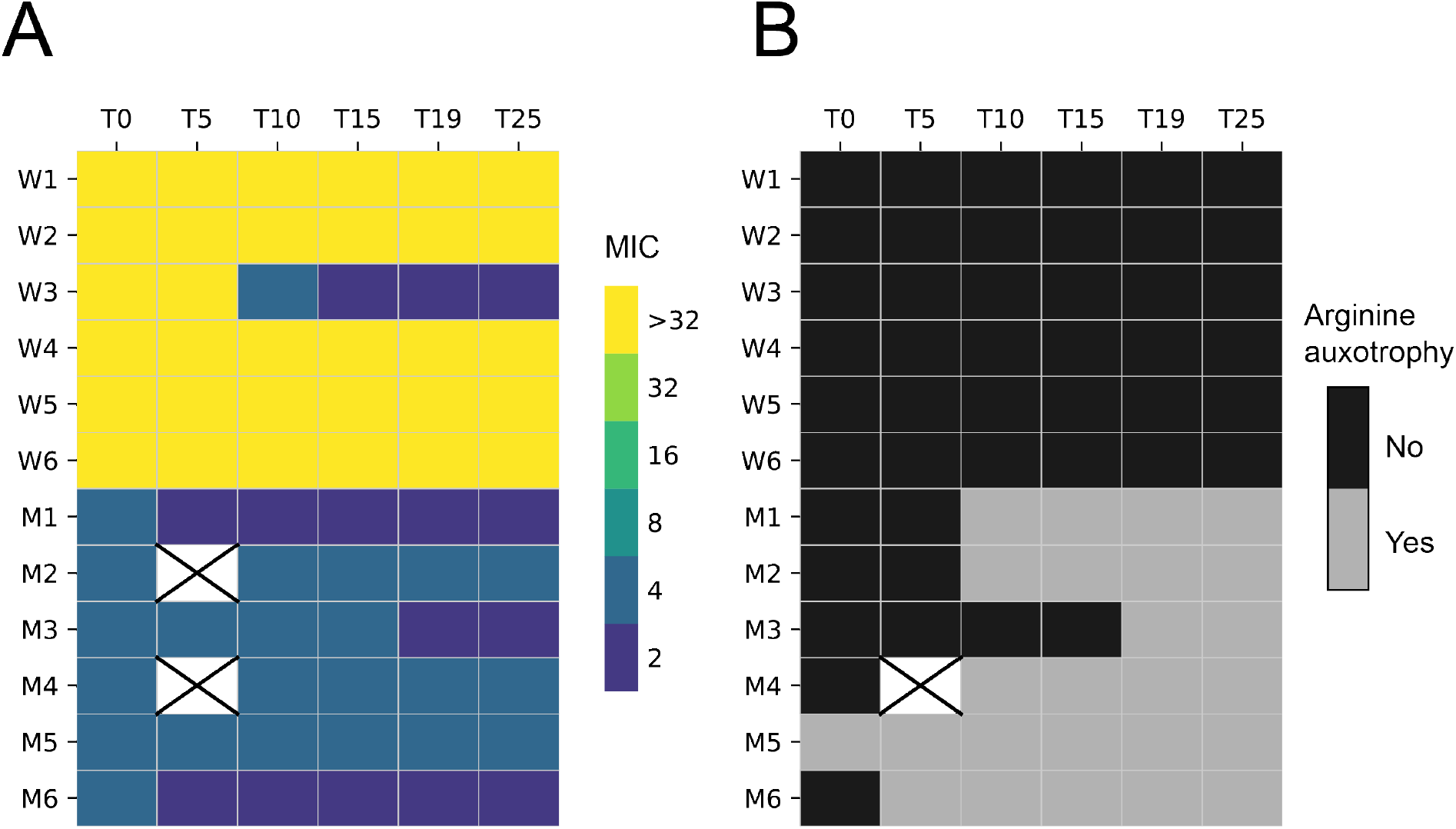
Phenotypic results for transferred lineages based on two genetic makers on the right chromosome arm. (**A**) MIC (μg ml^-1^) of chloramphenicol over time. (**B**) Arginine auxotrophy over time.

### Fitness and antibiotic production in evolved populations

Results in Fig. 3A show that the CFU of mutant lineages declined continuously compared to WT lineages. M2, that went extinct after the 5^th^ transfer, was only evaluated for the first two time points, and W3 was treated as a mutant lineage from the 7^th^ transfer, as explained above. Of the mutant lineages, all 7 showed significant reductions in CFU during the experiment (Welch’s *t* tests, all *P* < 0.01), amounting to a 9.8-fold median decline (IQR 5.4-13.3; one-sample Wilcoxon signed rank test, *P* = 0.016). By contrast, 4 of 6 WT lineages show small, but significant, increases in CFU (Welch’s *t* tests, all *P* < 0.05), amounting to a 2.4-fold median fitness increase (IQR 1.6-2.8; one-sample Wilcoxon signed rank test, *P* = 0.031). Accordingly, as shown in Fig S3, the median CFU change of WT and mutant lineages is significantly different from each other (Wilcoxon rank sum test, *P* = 0.0012).

**Fig. 3.**
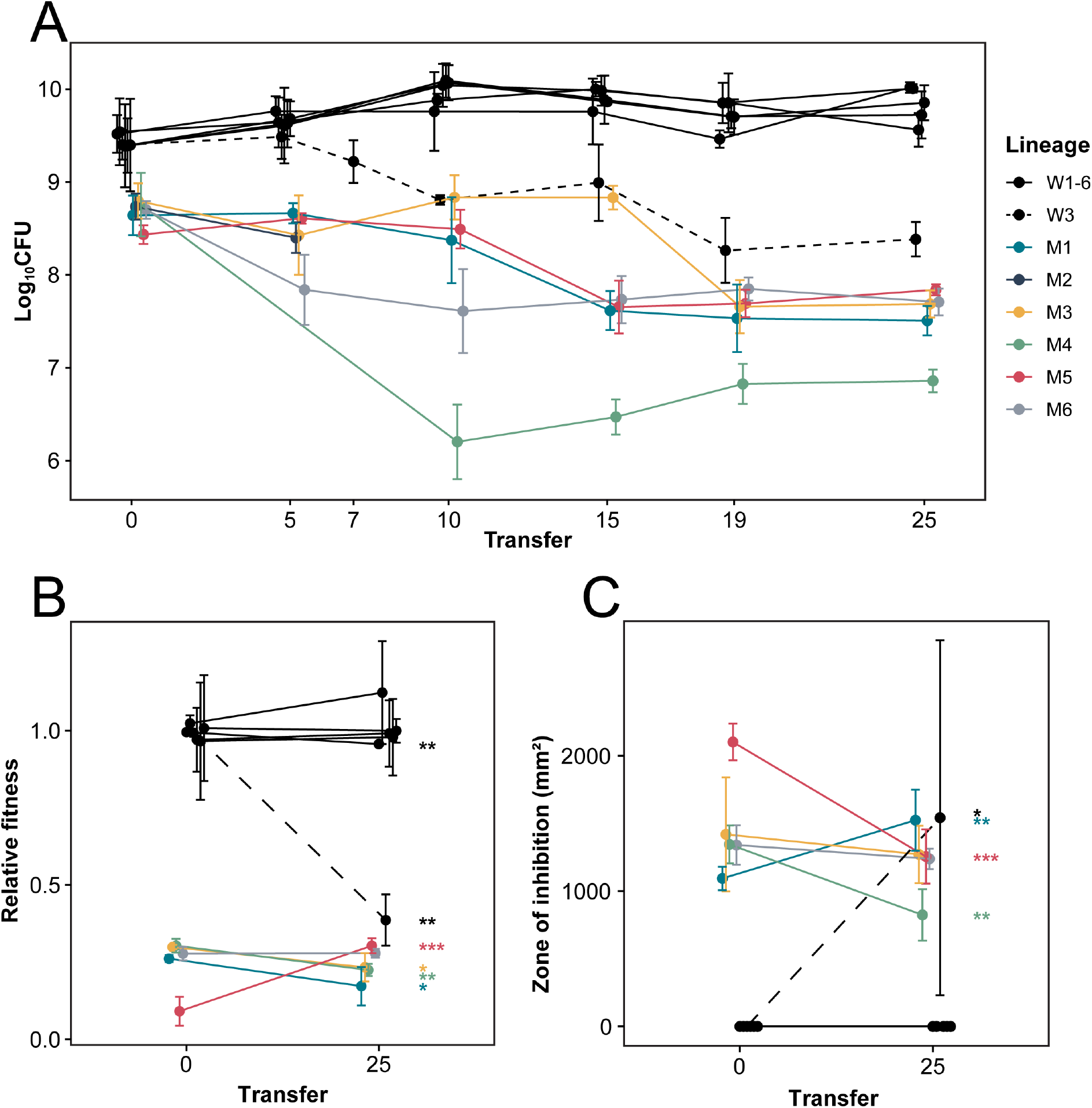
Fitness and antibiotic production in evolved lineages. (**A**) The fitness in terms of CFU of each lineage through time. WT lineages are shown in black with the one lineage W3 distinguished by a dashed line to indicate a conversion to mutant form after the 7^th^ transfer. The mutant lineages all show an overall decline through time with one lineage, M2, completely dying after the 5^th^ transfer. (**B**) Relative fitness of ancestor and evolved populations during competition with the WT. WT lineages show no significant change over time except for W1 and W3. The evolved strains in the mutant lineages show an overall decline in relative fitness except for M5. (**C**) Antibiotic production measured as the zone of inhibition for the ancestor and evolved strains of all lineages. WT lineages except for W3 show no change in antibiotic production. Mutant lineages show significant differences in antibiotic production. All error bars represent 95% confidence interval. Levels of statistical significance are indicated as * (< 0.05), **(<0.01) and ***(<0.001) (Welch’s *t* test).

To further examine changes during mutation accumulation, the relative fitness of all ancestral and evolved populations were measured using pairwise competition assays. Results in Fig. 3B are consistent with those based on CFU, and the two fitness measures are significantly correlated (F_1,20_ = 59.93, r^2^ = 0.75, *P* = 1.928×10^-7^) (Fig. S4). Except for W1 and W3, the fitness of WT lineages remain unchanged, while most mutant lineages have significantly reduced fitness (Welch’s *t* tests, all *P* < 0.05). The fitness of one lineage (M5) surprisingly increased from its initial value, suggesting that CFU, which measures sporulation rate, may also be impacted by other aspects of *Streptomyces* multicellular growth that are important during fitness assays.

Finally, we estimated antibiotic production of evolved lineages using an inhibition assay (Fig. 3C). As expected, mutant strains produce significantly larger halos than WT strains at the start (Wilcoxon rank sum test with continuity correction, *P* = 0.00389) and end (Wilcoxon rank sun test with continuity correction, *P* = 0.00549) of serial transfer. While WT strains remain unchanged, and produce small zones of inhibition against the *B. subtilis* target species, average antibiotic production in all mutant strains remains extremely high, while three lineages show small, but significant, shifts (Welch’s *t* tests, all *P* < 0.01). Notably, antibiotic production in W3 increased markedly, coincident with its decreased fitness; this change confirms the strong correlation between these two traits ^13^.

### Continuous deletions in mutant lineages but not wild-type lineages

To identify genetic changes that led to the rapid declines in mutant CFU, we used whole-genome sequencing to measure changes in genome size by mapping against a reference strain (Fig. S5). As expected, no changes were observed in WT lineages (with the exception of W3 from the 7^th^ transfer). By contrast, as shown in Fig. 4A and Fig. S5, mutant lineages continued to accumulate large deletions to the left and right chromosome arms during serial transfer. Deletions to the left arm ranged from 0 to 882 kb, and in the right arm from 0 to 250 kb (Left arm: 289 ± 117 kb (mean ± SE), n = 7; Right arm: 80 ± 30 kb (mean ± SE), n = 7). The total deletion size of these strains ranged from 0 to 924 kb (369 ± 124 kb (mean ± SE), n = 7). One lineage (M2) suffered an abnormally large deletion on the left chromosome arm, and this strain was no longer able to develop an aerial mycelium, resulting in a bald phenotype (Fig. S1). However, no apparent deletions in known *bld* genes could be identified ^24^, suggesting other causes for this phenotype. Additionally, one lineage (M5) that began with the shortest genome did not gain further deletions, suggesting that further genome loss may not have been possible due the presence of essential genes near to the border of the chromosome ends. Intriguingly, this is also the only mutant lineage in which we observed an increased relative fitness during the MA experiment (Fig. 3B), suggesting that changes to this strain were due to point mutations. Fig. 4B plots the relationship between CFU and the sizes of genomic deletions on the left arm, right arm or entire chromosome. These results confirm and extend our previous observations. CFU and deletion size are negatively correlated for the left arm (F_1,11_ = 6.026, r^2^ = 0.354, *P* = 0.032), the right arm (F_1,11_ = 9.881, r^2^ = 0.473, *P* = 0.009) and for the whole chromosome (F_1,11_ = 10.75, r^2^ = 0.494, *P* = 0.007).

**Fig. 4.**
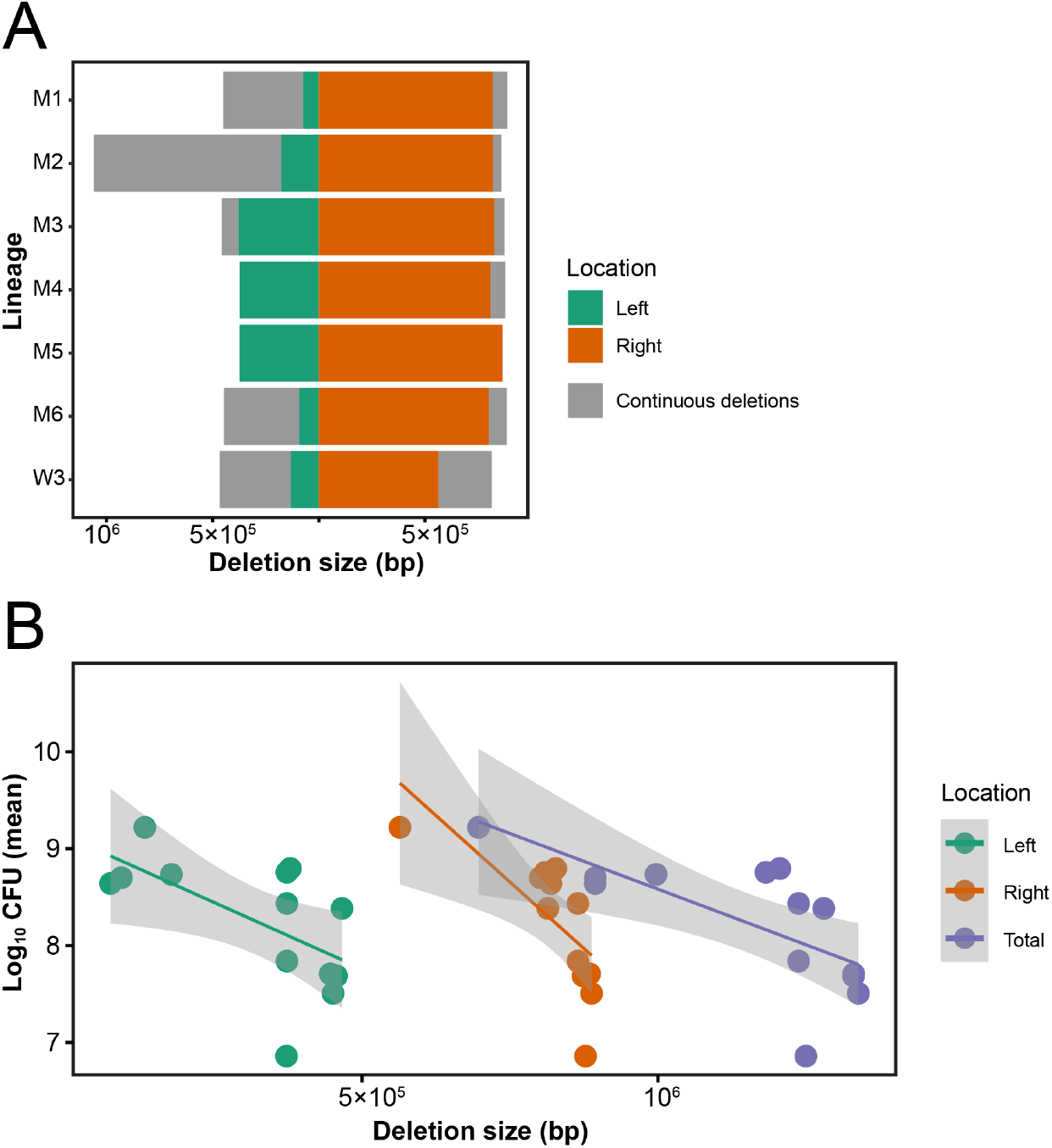
Genomic deletions and their effects on strain fitness. (**A**) Initial and final deletion sizes on the left and right chromosome arms. Green and orange regions represent deletions on the left and right chromosome arms of ancestral strains, while regions in gray represent new deletions in evolved strains. Sequenced strains include M1-M6 and W3 at T0 (or T5 for W3) and T25. (**B**) Significant negative correlation between the size of the chromosome deletions and strain fitness, shown for the left arm, the right arm and the entire genome. Statistics are given in the main text.

### Increased base-substitution rates in mutant lineages

To address other sources of mutational variation, in addition to gross chromosome changes, we estimated the base-substitution and indel mutation rates from mutant and WT lineages. Unexpectedly, we found that mutant lineages fixed significantly more mutations than the WT lineages. Overall, mutants fixed 29.5 mutations per lineage (median, IQR 12.25-32.5, n = 6) while the WT lineages fixed 5 mutations per lineage (median, IQR 4-6, n = 5). To account for differences in the number of transfers of different lineages (due to the impact of W3 that became a mutant after 5^th^ transfer), we calculated a per transfer mutation rate. This analysis showed that the base-substitution rate for mutants was 12.78 per 10^8^ nucleotides per transfer (median, IQR 7.62-17.46, n = 7) compared to 1.5 per 10^8^ nucleotides per transfer (median, IQR 1.28-2.03, n = 6) in WT, exhibiting a roughly 10-fold difference (Wilcoxon rank sum test with continuity correction, *P* = 0.018) (Fig. 5A). When we partitioned this result into different mutant classes, we observed that mutants acquired synonymous and non-synonymous mutations as well as changes in non-coding regions at a significantly higher rate (Fig. 5B). Further, looking across different transitions and transversions, we found that mutants fixed more mutations in 4 out of 6 mutation classes (Fig. 5C). Four mutant lineages fixed mutations in alleles affecting DNA replication or repair ^19,20^, including DNA polymerase III (synonymous), DNA topoisomerase IV (synonymous), DNA polymerase I (non-synonymous) and DNA ligase (non-synonymous) (Tables S1 and S2). Although suggestive, at present we cannot confirm that these specific changes are causally associated with increased mutation fixation.

**Fig. 5.**
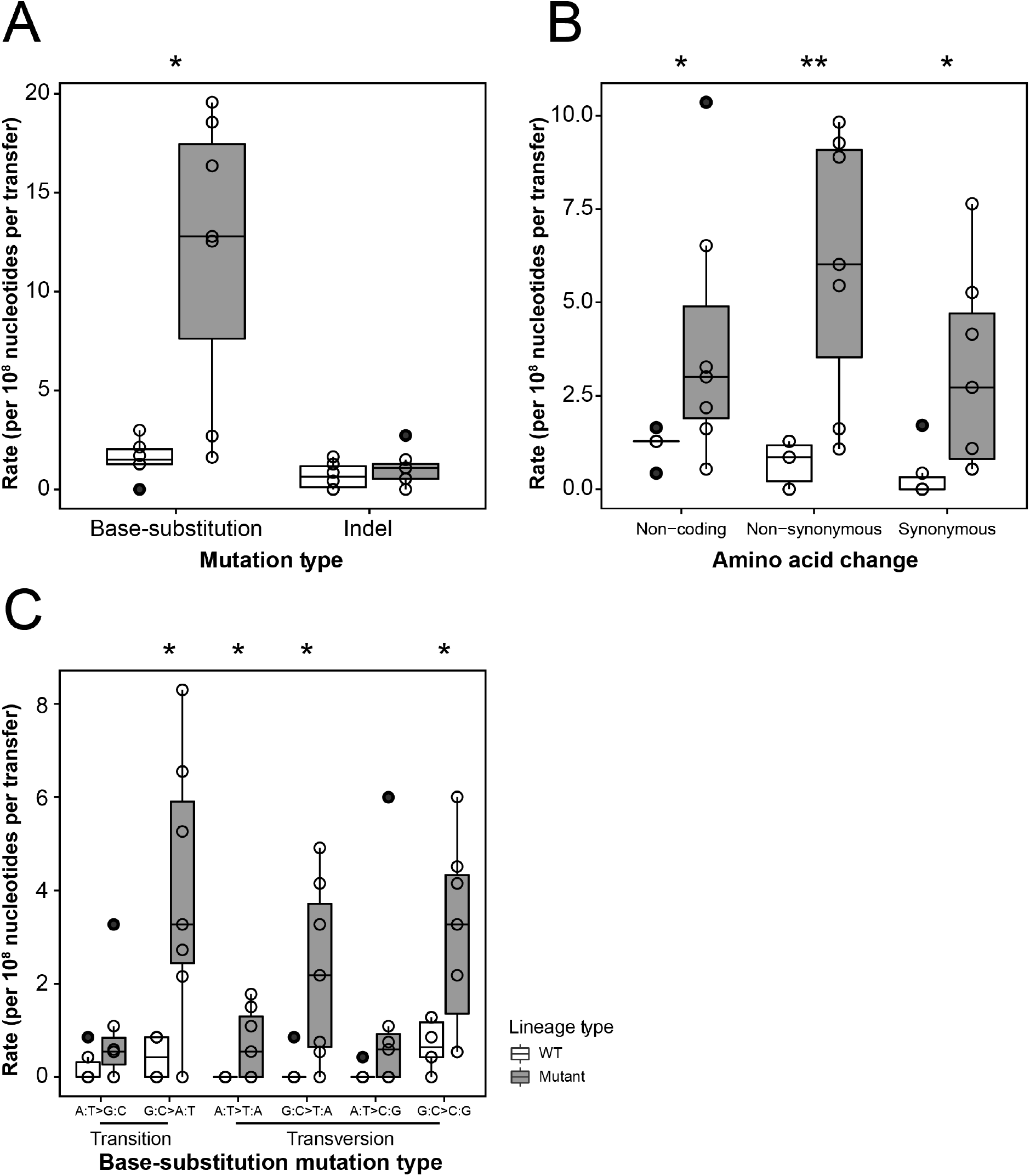
Mutation rates of WT and mutant lineages for different mutation classes. Mutation rates are partitioned according to: (**A**) Base-substitutions and indels; (**B**) the types of amino acid changes; and (**C**) for different classes of transitions or transversions. Levels of statistical significance are indicated as * (*P* < 0.05) and ** (*P* < 0.01) (Wilcoxon rank sum test).

These results thus indicate that mutant lineages become mutators, in addition to acquiring large genomic deletions. Both factors likely contribute to their dramatic fitness reductions.

## Discussion

Division of labor allows populations of individuals to more efficiently carry out functions that are mutually incompatible ^3,25,26^. In microbes, division of labor can facilitate biofilm formation ^25,27^, energy transfer^28^, and coordinated metabolism ^29^, among other behaviors. In some cases, division of labor leads to sub-populations of cells that carry out functions that are lethal to themselves but that benefit the entire colony ^30^. For example, colicin secretion in *E. coli* requires cell lysis ^31^, a fate limited to a small fraction of cells with low reproductive value. By this process, the burden of colicin-secretion is disproportionately borne by the cells with the least to lose in terms of their own fitness ^32,33^. We recently provided evidence for a similar phenomenon in *Streptomyces*, whereby a subfraction of cells within a multicellular colony hyperproduces antibiotics at the expense of their own reproduction, in part due to large and irreversible deletions from their chromosome ends ^13^. The aim of the present work was to examine the fate of these altruistic cells after their emergence. We found that although *Streptomyces* cells hyper-producing antibiotics do not lyse, like *E. coli* colicin producers, they continue to accumulate large deletions and also evolve an increased mutation rate across their genome. These effects, which lead to an “effective lethality”, suggest that these cells are analogous to the reproductively sterile worker castes in social insects ^34^.

Our experimental approach was designed to approximate the natural growth and development of *Streptomyces* that disperse via spores, such that each new colony passes through a single-cell bottleneck. This resembled a classic mutation accumulation experimental design, which has been widely used to examine fitness declines in microbes due to the accumulation of deleterious mutations via Muller’s Ratchet, a process in which deleterious mutations accumulate irreversibly in a population lacking recombination ^35,36^. As in mutation accumulation experiments, our mutants lost fitness ^17,18^; however, their rate of decline was more rapid due to mutations of very large effects via genome loss as well as point mutations. Results in Fig. 4B show a significant negative relationship between total genome size and CFU production, consistent with studies performed in *E. coli* containing manipulated reduced genomes ^37^. Given the 679-1817 genes that are lost from these populations, it is not possible to know which ones are responsible for the fitness reductions, either alone or in combination. In addition to genome loss, we were surprised that mutant lineages, but not wild-type ones, have an approximately 10-fold increased mutation rate, likely due to mutations in genes for DNA replication and repair^19,20^. Mutations were found in several mutation classes and are higher in both coding and non-coding regions, indicating broad and non-specific mutagenesis. Decreased competitiveness, massively compromised CFU as cells pass through single-spore bottlenecks, and the combined accumulation of large deletions and an increased mutation burden, lead to synergistic declines in fitness that resembles a type of mutational meltdown. First, ecologically deficient mutants develop a higher mutation rate. Second, these lineages accumulate further deletions, which magnifies their fitness reductions and causes an irreversible decrease in their effective population size, ultimately leading to extinction. Although this process occurs within an organism over a very short time period, it closely resembles the idea of a classical mutational meltdown, in which a small population going through Muller’s ratchet experiences accelerating fitness declines caused by deleterious mutations ^38^.

Even though mutant lineages are deteriorating at a pace that exceeds results from other MA experiments, they don’t die immediately, as do *E. coli* colicin producers. Why do antibiotic producing strains of *Streptomyces* die via mutational meltdown instead of lysing? One possible cause of this difference may be the intrinsic differences in the activity of antibiotics and colicins. Whereas the latter can act at very low concentrations, *e.g.* via single-hit kinetics ^31^, antibiotics may require higher concentrations to provide sufficient protection to large *Streptomyces* colonies. Antibiotics can also bind tightly to abiotic substrates, potentially requiring higher levels of production within colonies ^39^. These possibilities would necessitate continued survival and growth of producing cells, thereby generating spatially clustered mutant sub-populations within colonies that hyperproduce antibiotics, whereas sufficient toxin quantities could be produced by single *E. coli* cells either dispersed randomly throughout the colony or on the colony edge facing impending threats ^32,33^. A related issue that remains unresolved is the origin of mutant cells within growing colonies. Specifically, it remains unclear if low-level antibiotic production somehow causes subsequent genome decay due to local toxicity, if stochastically damaged cells subsequently adopt a new fate to hyper-produce antibiotics, or if mutants and the fraction of cells that adopt this fate are induced by local competitive conditions in the soil environment. At present, we are unable to fully address these issues and they remain important areas for future study.

Altruistic behaviors can be explained by their indirect fitness benefits, whereby individuals offset the loss of their own reproduction by increasing the reproduction of their relatives ^40^. In multicellular bacteria, like streptomycetes or cyanobacteria, clonality, and therefore high relatedness, among cells in the colony is ensured by their mode of filamentous growth ^9,10^. For this reason, division of labor with extreme specialization can evolve and lead to the elaboration of multiple cell types. Streptomycetes are typically divided into two functional classes of cells: spores and vegetative cells ^11,12^. Our work supports the notion that colonies can be further partitioned into at least one more cell type, those producing antibiotics and that accumulate extreme and irreversible genetic damage leading to their demise. We would predict similar diversification among other streptomycetes, as well as the discovery of additional divisions of labor among other multicellular bacteria ^9^.

## Methods

### Bacterial strains and cultural conditions

Strains used in this study all derived from *Streptomyces coelicolor* A3 (2) M145. Strains were maintained and assayed at 30°C on soy flour mannitol media (SFM) containing 20 g mannitol, 20 g agar and 20 g soya flour per liter of water. Spores of *S. coelicolor* were diluted and plated onto an SFM plate. To obtain initial isolates for mutation accumulation, one random WT colony (designated as WT_ancestor_) was diluted and plated onto SFM agar. One random WT and mutant colony were then picked and replated onto separate SFM plates. Six random colonies were then chosen from each plate and designated as ancestors for subsequent serial passage through single-colony transfer, for a total of 12 lineages (6 WT and 6 mutants). During each transfer, a single colony from each lineage growing closest to a randomly placed spot on the back of the plate was chosen and streaked onto another SFM plate. This procedure was repeated every 3-4 days for 25 transfer cycles (Fig. 1B). Transferred lineages were archived by creating a full lawn from the transferred colony, after which spores were harvested after ~ 7 days of growth and sporulation as previously described ^41^. All stocks were maintained at −20°C.

### Competition assay

We estimated the fitness of mutant (M1, M3, M4, M5 and M6) and WT lineages (W1-6) from TO and T25 (n = 2 per WT strain and n = 4 per mutant strain), following the protocol in ^13^. All strains were marked with apramycin resistance and the WT ancestor was marked with hygromycin B resistance, by using integrating plasmids pSET152 and pIJ82, respectively. After diluting strains to 10^6^ CFU ml^-1^, they were mixed with the reciprocally marked WT ancestor at different initial frequencies. 100 μl of each mixture was plated onto 25 ml SFM agar plates and incubated at 30°C for 5 days. At the same time, each mixture was serially diluted and plated onto SFM agar plates containing either apramycin (50 μg ml^-1^) or hygromycin B (50 μg ml^-1^) to obtain precise estimates of initial frequencies. After 5 days, each plate was harvested in H_2_O and passed through an 18-gauge syringe plugged with cotton wool to remove mycelial fragments and resuspended in 1 ml 20% glycerol. Each sample was then serially diluted onto plates containing either antibiotic to calculate final frequencies.

### Antibiotic production

All mutant and wildtype populations from T0 and T25 were tested for antibiotic production by measuring the zone of inhibiton in an overlay assay. Spores were diluted to get 10^6^ spores ml^-1^ and subsequently 2 μl was spotted onto an SFM plate in the center. Three replicates of each strain were prepared and incubated for 5 days at 30°C. On the 6^th^ day, exponentially growing cultures of *B. subtilis* were prepared by subcultring an overnight grown strain in LB medium and incubated until OD_600_ 0.4-0.6 was achieved. Soft agar was prepared by mixing LB liquid with LB agar in a 1:1 ratio and maintained at 60°C. 150 μl of the exponentially growing *B. subtilis* was added to 7 ml of soft agar, mixed adequately and poured onto the SFM plate containing the *Streptomyces* spots growing in the center. The soft agar was allowed to solidify and the plates were subsequently incubated overnight at 30°C. The plates were scanned the next day and the area of inhibition/clearance was measured using the ImageJ/Fiji software.

### Estimating antibiotic resistance

To estimate changes to antibiotic resistance, minimal inhibitory concentration (MIC) was determined for all strains by spot dilution onto large SFM agar plates (150 x 20 mm, Sarstedt, Germany) supplemented with different antibiotic concentrations. Drug concentrations ranged from 2 to 32 μg ml^-1^ (chloramphenicol, oxytetracycline and ciprofloxacin) and 1 to 16 μg ml^-1^ (streptomycin). Plates were inoculated using a 96-pin replicator from master 96-well plates containing ~ 10^7^ spores ml^-1^. Approximately 1 μl from this stock was applied to each plate; the replicator was flame sterilized between each transfer to ensure that no cells or antibiotics were transferred between assay plates. The plates were incubated for 4 days at 30°C and then imaged and scored for growth. The MIC was determined as the drug concentration where no growth was visible after 4 days (n = 3 per strain per drug concentration).

### Auxotrophy assay

To test for auxotrophy, strains were grown on minimal media (MM) containing per liter 0.5 g asparagine, 0.5 g K_2_HPO_4_, 0.2 g MgSO_4_·7H_2_O, 0.01 g FeSO_4_·H_2_O and 10 g agar, supplied with either 0.5% mannitol or 0.5% mannitol plus 0.0079% arginine. Bacteria were spotted onto plates using a pin-replicator, as for MIC assays, and grown for 4 days at 30°C. Auxotrophy was detected by comparing growth of colonies on plates with or without supplemented arginine (n = 3 per strain).

### CFU estimation

We used CFU to estimate the fitness of strains from each lineage. For each strain, 10^5^ spores were plated onto SFM as a confluent lawn. After 5 days of growth, spores were harvested by adding 10 ml H_2_O to the plates, gently scraping the plate surface to create a spore suspension, and then filtering the liquid through an 18-gauge syringe with cotton wool to remove mycelia. After centrifugation, spore stocks were resuspended in 1 ml 20% glycerol and then serially diluted onto SFM to calculate the total CFU for each strain (n = 3 per strain, except n = 2 for M1 at T19).

### Whole-genome sequencing

Sequenced strains include WT_ancestor_, M1-M6 at T0, W3 at T5, W3 at T7, and all lineages at T25. Strains were sequenced using two approaches. Long reads sequencing (PacBio, USA) was performed as previously reported ^13^. Short reads sequencing (BGISEQ-500) was done using the following protocol. DNA was extracted after growth in liquid TSBS: YEME (1:1 v:v) supplemented with 0.5% glycine and 5 mM MgCl_2_. Approximately 10^8^ spores were inoculated in 25 ml and incubated at 30°C with a shaking speed of 200 rpm for 12-48 hours. TSBS contains 30 g tryptic soya broth powder and 100 g sucrose per liter and YEME contains 3 g yeast extract, 5 g peptone, 3 g malt extract, 10 g glucose and 340 g sucrose per liter. DNA was extracted using phenol/chloroform ^41^. Visible cell pellets were washed with 10.3% sucrose solution after centrifugation. Pellets were resuspended in 300 μl GTE buffer, containing 50 mM glucose, 10 mM EDTA, 20 mM Tris-HCl, pH 7.5 and 4 mg ml^-1^ lysozyme and incubated at 37°C for 1 hour. Then 300 μl 2M NaCl was added and gently inverted ten times, followed by the addition of 500 μl phenol/chloroform (bottom layer). After mixing, each tube was centrifuged for 5min and the upper layer was transferred to a new tube. This procedure was repeated at least twice until the intermediate layer was almost invisible. The final transferred upper layer was mixed with a same volume of 2-proponol, and centrifugated for 10 min. Liquid in the supernatant was discarded and pellets were dried at room temperature before being dissolved in 200 μl Milli-Q H_2_O. After adding 1 μl RNase, the DNA was resuspended at 37°C for 1 hour. Phenol/chloroform washing and DNA precipitation was repeated once to remove the RNase. After adding phenol/chloroform, the upper layer was transferred to a new tube, and then mixed with 16 μl 3M pH 5.2 NaCH_3_COO and 400 μl 96% ethanol. This mixture was cooled at −20°C for 1 hour and centrifuged for 10 min to obtain the DNA pellets. Pellets were washed with pre-cooled 96% ethanol and dried at room temperature. DNA was dissolved in Milli-Q H_2_O and sent for commercial sequencing at BGI (Hong Kong).

### Sequencing processing

The raw data of PacBio sequencing was processed as outlined in Zhang *et al.* (2020) ^13^ and genome length was evaluated based on these results. The BGISEQ-500 data was handled using CLC Genomics Workbench (QIAGEN, v 8.5.4). Filtered raw reads were first imported and mapped to the reference genome NC_003888.3 ^42^ through the “NGS core”- “Map to the reference” function. Variants were then called by using “Basic variant detection” function, with the filter parameters set to minimum coverage as 5, minimum count as 2 and minimum frequency as 50%. Variants were identified by comparing lineages to their corresponding parental strain by applying the “Resequencing analysis” - “Compare Variants” - “Compare Sample Variant Tracks” option. By using the annotation information in the GenBank file, final variants were then annotated by applying the “Track tools” - “Annotate with overlap information” option, and amino acid changes were added to the variant track by “Resequencing analysis”- “Functional consequences” - “Amino acid changes” option. Final results were exported as excel sheets and variants in genes that were not detected in PacBio sequencing were removed before performing further analyses.

### Statistical analyses

All statistical analyses were performed in R (v 3.6.2). Welch’s *t* test was used to test differences of CFU production, relative fitness and antibiotic production across the course of the experiment, following the Shapiro-Wilk normality test. One-sample Wilcoxon signed rank test was used to test if the CFU change after transfers while Wilcoxon rank sum test was used to compare the difference between WT and mutant lineages. All tests are two-sided.

## Data availability

All data will be provided along with the published version of this article

## Author contributions

Z.Z., D.C. and D.E.R conceptualized this study. Z.Z., S.S. and B.C. performed the experimental work. Z.Z., S.S., D.C. and D.E.R. analyzed the results. Z.Z., S.S., D.C. and D.E.R. wrote the manuscript. All authors approved the final submitted version of this article.

## Acknowledgements

We would like to thank Chao Du for his help with bioinformatics and data visualization.

