## Supplemental figures for "Mutational meltdown of microbial altruists in *Streptomyces coelicolor* colonies"

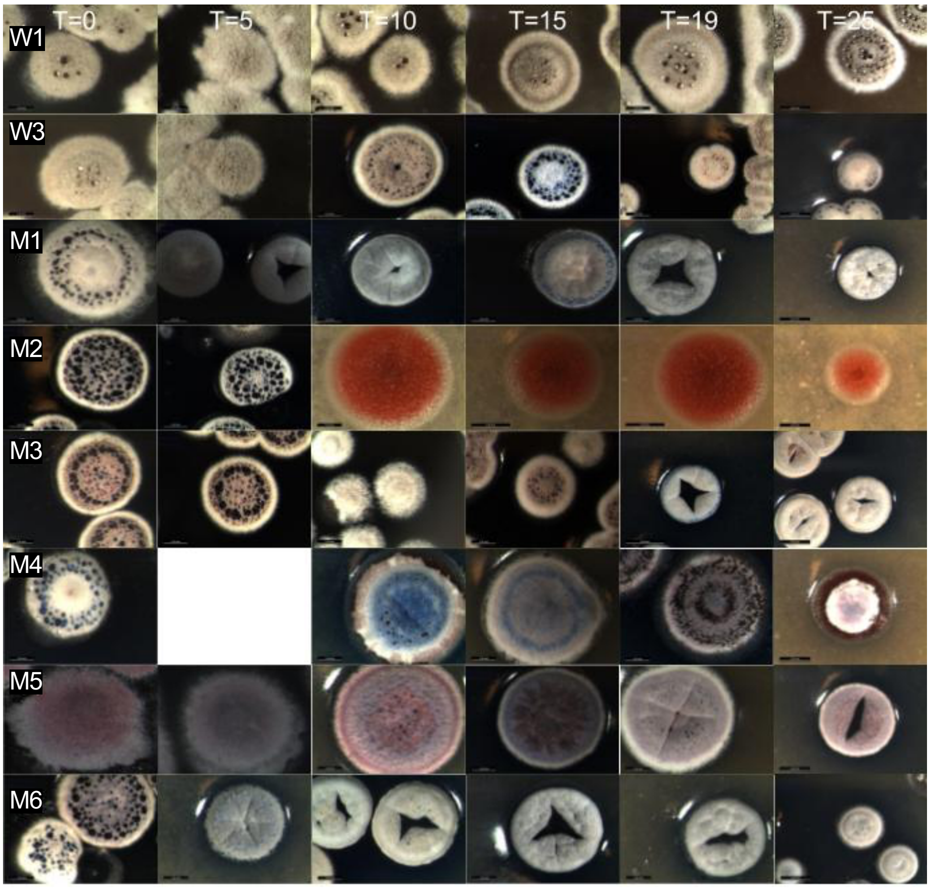


**Fig. S1. Morphology of sampled strains.** Morphogenesis, including aerial growth, sporulation and pigmentation, varies among and within lineages.

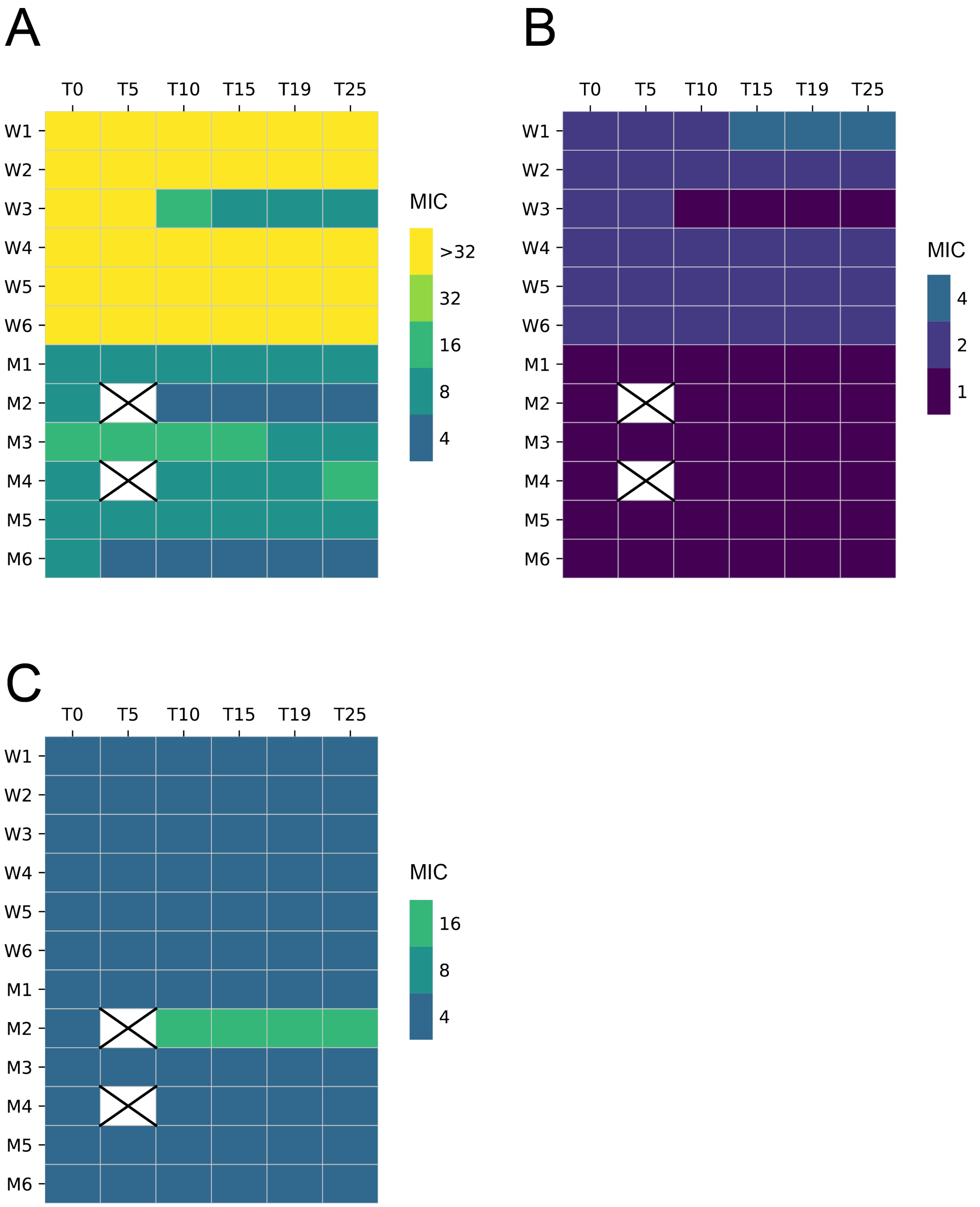


**Fig. S2. Antibiotic resistance of sampled strains.** MIC of (**A**) oxytetracycline (**B**) streptomycin or (**C**) ciprofloxacin. Unit is shown as μg $ml^{-1}$.

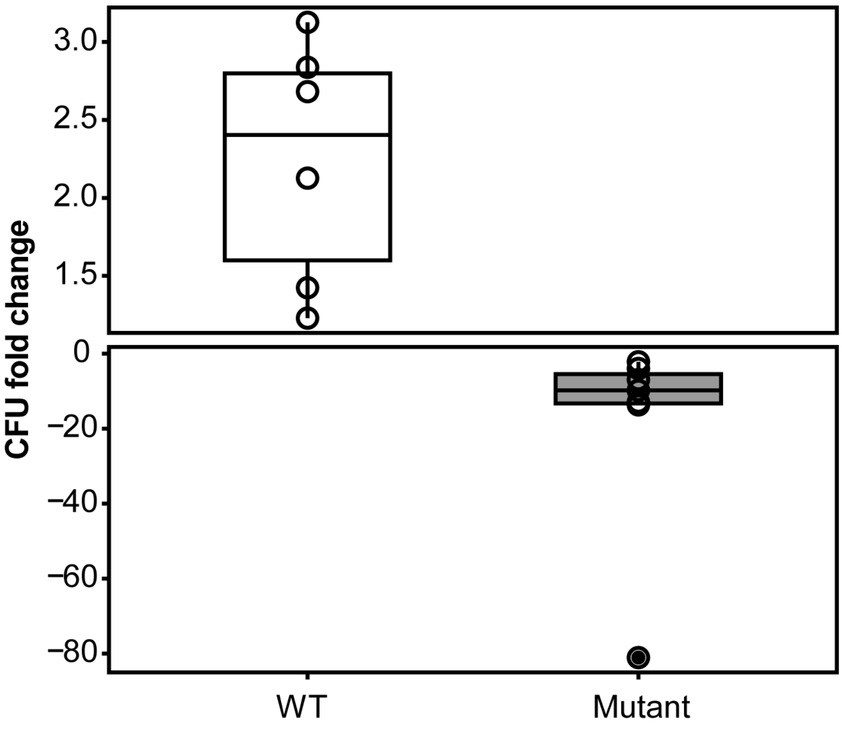


**Fig. S3. The fold change of CFU during the MA experiment.** The median fold changes of CFU of WT and mutant lineages are shown.

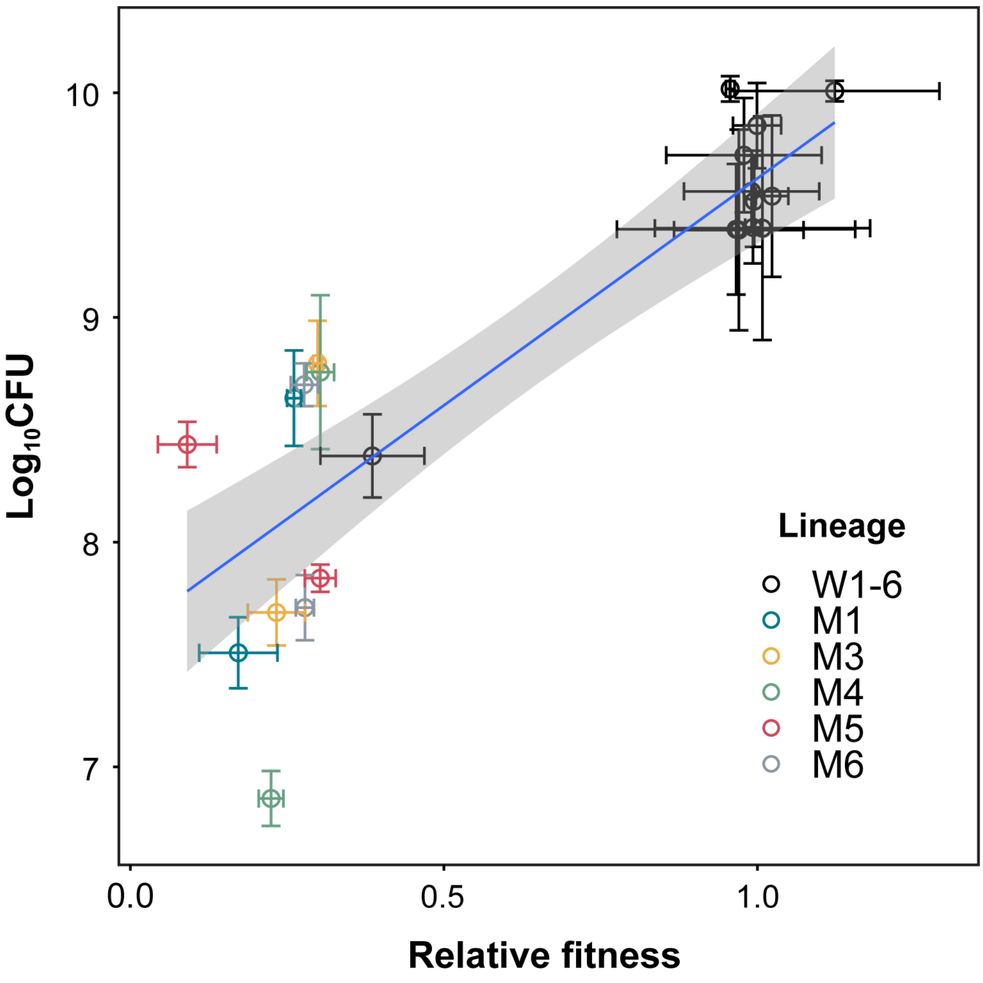


**Fig. S4. Correlation between relative fitness and CFU.** A significant correlation is observed for sampled ancestral and evolved strains from different lineages.

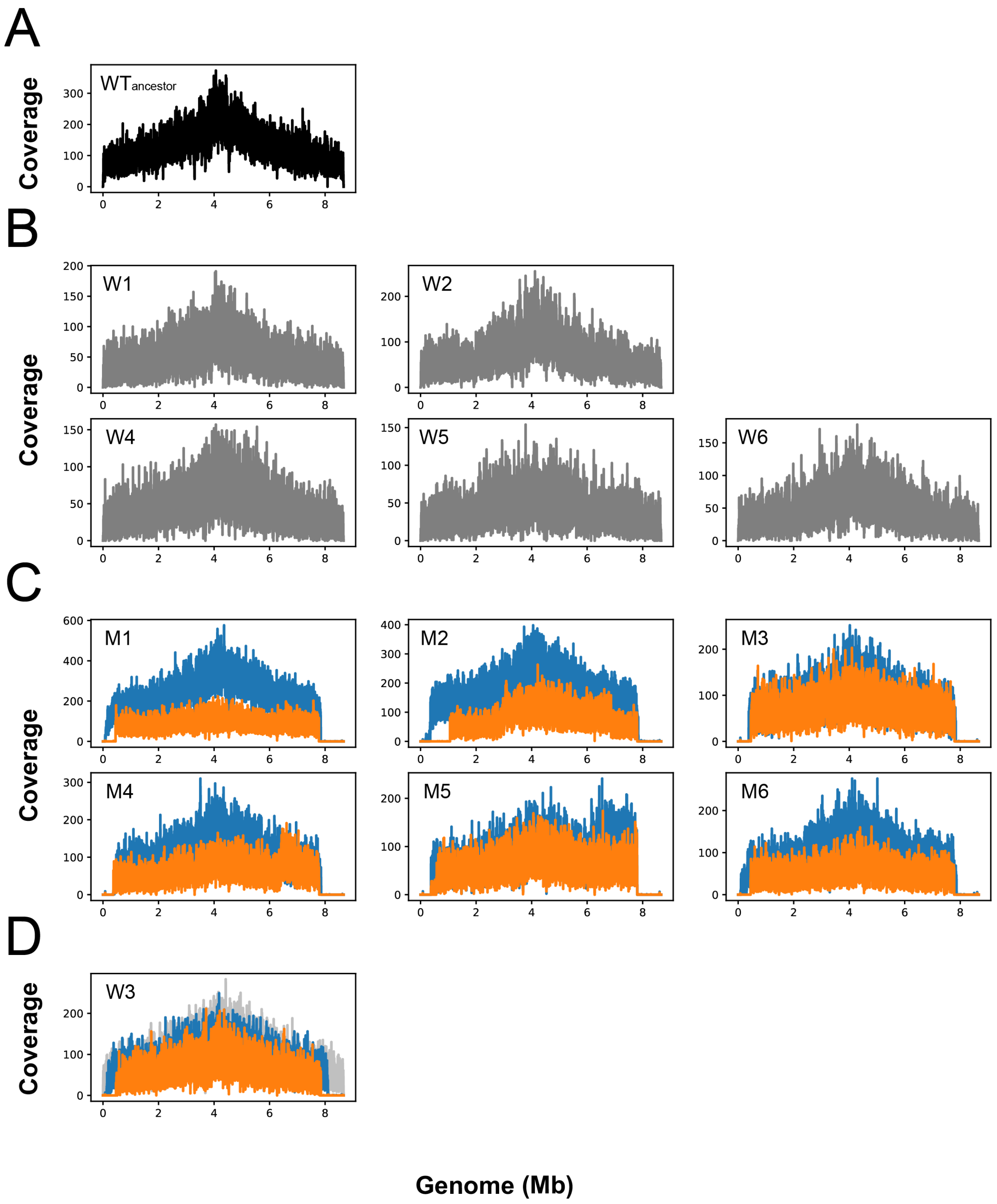


**Fig. S5. Pacbio sequencing results of sampled strains.** Plots indicate the coverage of reads mapped to the *S. coelicolor* M145 reference genome. (**A**) Ancestor WT. (**B**) T25 strains from WT lineages. (**C**) T0 (blue) and T25 (orange) strains from mutant lineages. (**D**) Strains of lineage W3 at T5 (light gray), T7 (blue) and T25 (orange).
