## Supplemental table S1 for "Mutational meltdown of microbial altruists in *Streptomyces coelicolor* colonies"

### Table S1. Accumulated mutations in wild-type lineages during the experiment.

| Lineage | Region | Type | Reference | Allele | Count | Coverage | Frequency | Coding region change | Amino acid change | Non-synonymous | Mutation type | Gene | Gene product |
|---|---|---|---|---|---|---|---|---|---|---|---|---|---|
| | 221017 | SNV | C | G | 46 | 73 | 63.0137 | | | - | G:C>C:G | | |
| | 2208542 | SNV | G | A | 122 | 124 | 98.3871 | NP_626320.1:c.272C>T | NP_626320.1:p.Ala91Val | Yes | G:C>A:T | SCO2060 | integral membrane transport protein |
| | 2349467 | SNV | A | C | 166 | 168 | 98.80952 | NP_626436.1:c.728A>C | NP_626436.1:p.Glu243Ala | Yes | A:T>C:G | SCO2183 | 2-oxoacid dehydrogenase subunit E1 |
| | 2462887..246288 | Deletion | GG | - | 100 | 161 | 62.1118 | | | - | Deletion | rnpB | |
| | 4148255 | SNV | G | C | 15 | 30 | 50 | | | - | G:C>C:G | | |
| W1 | 8355042 | SNV | C | T | 87 | 87 | 100 | NP_631579.1:c.308C>T | NP_631579.1:p.Ser103Leu | Yes | G:C>A:T | SCO7535 | lipoprotein |
| | 221017 | SNV | C | G | 35 | 61 | 57.37705 | | | - | G:C>C:G | | |
| | 4744885 | SNV | C | G | 199 | 204 | 97.54902 | NP_628502.1:c.2109C>G | NP_628502.1:p.Asp703Glu | Yes | G:C>C:G | SCO4331 | hypothetical protein |
| | 5939426 | SNV | G | C | 118 | 119 | 99.15966 | NP_629591.1:c.152G>C | NP_629591.1:p.Gly51Ala | Yes | G:C>C:G | SCO5454 | two-component system sensor kinase |
| | 6176177^617617 | Insertion | - | G | 52 | 96 | 54.16667 | | | - | Insertion | | |
| W2 | 7313624 | Deletion | C | - | 21 | 39 | 53.84615 | | | - | Deletion | | |
| W3 | 2458948^245894 | Insertion | - | C | 69 | 107 | 64.48598 | | | - | Insertion | | |
| | 221017 | SNV | C | G | 41 | 73 | 56.16438 | | | - | G:C>C:G | | |
| | 3739268 | SNV | G | A | 226 | 226 | 100 | | | - | G:C>A:T | | |
| | 7988760 | SNV | A | G | 57 | 57 | 100 | | | - | A:T>G:C | | |
| W4 | 8613917 | SNV | C | T | 70 | 70 | 100 | NP_631818.1:c.129G>A | | No | G:C>A:T | SCO7788 | hypothetical protein |
| | 373226 | SNV | G | C | 5 | 9 | 55.55556 | NP_624691.1:c.182G>C | NP_624691.1:p.Ser61Thr | Yes | G:C>C:G | SCO0368 | transposase |
| | 1650273 | SNV | A | G | 127 | 128 | 99.21875 | | | - | A:T>G:C | | |
| | 2458948^245894 | Insertion | - | C | 75 | 126 | 59.52381 | | | - | Insertion | | |
| | 2462887 | Deletion | G | - | 75 | 124 | 60.48387 | | | - | Deletion | rnpB | |
| | 4938336 | SNV | C | A | 308 | 309 | 99.67638 | NP_628680.1:c.185G>T | NP_628680.1:p.Arg62Leu | Yes | G:C>T:A | SCO4516 | hypothetical protein |
| | 5371295 | SNV | G | T | 167 | 168 | 99.40476 | NP_629088.1:c.1227G>T | | No | G:C>T:A | SCO4935 | hypothetical protein |
| | 8424413 | SNV | G | A | 18 | 18 | 100 | NP_631639.1:c.465C>T | | No | G:C>A:T | SCO7597 | hypothetical protein |
| | 8424416 | SNV | G | A | 17 | 17 | 100 | NP_631639.1:c.462C>T | | No | G:C>A:T | SCO7597 | hypothetical protein |
| | 8424506 | SNV | A | G | 19 | 19 | 100 | NP_631639.1:c.372T>C | | No | A:T>G:C | SCO7597 | hypothetical protein |
| W5 | 8424513 | Deletion | G | - | 26 | 26 | 100 | NP_631639.1:c.365delC | NP_631639.1:p.Ala122fs | Yes | Deletion | SCO7597 | hypothetical protein |
| | 221017 | SNV | C | G | 51 | 77 | 66.23377 | | | - | G:C>C:G | | |
| | 6769943 | SNV | G | C | 15 | 28 | 53.57143 | NP_630271.1:c.536G>C | NP_630271.1:p.Arg179Pro | Yes | G:C>C:G | SCO6167 | proline rich protein membrane protein |
| W6 | 7700826 | SNV | C | G | 67 | 68 | 98.52941 | NP_631001.1:c.553G>C | NP_631001.1:p.Val185Leu | Yes | G:C>C:G | SCO6935 | hypothetical protein |
