## Supplemental table S2 for "Mutational meltdown of microbial altruists in *Streptomyces coelicolor* colonies"

### Table S2. Accumulated mutations in mutant lineages during the experiment.

| Lineage | Region | Type | Reference | Allele | Count | Coverage | Frequency | Coding region change | Amino acid change | Non-synonymous | Mutation type | Gene | Gene product |
|---|---|---|---|---|---|---|---|---|---|---|---|---|---|
| | 481270 | SNV | G | A | 239 | 241 | 99.17012 | NP_624779.1:c.259G>A | NP_624779.1:p.Ala87Thr | Yes | G:C>A:T | SCO0459 | hypothetical protein |
| | 712725 | SNV | C | T | 186 | 188 | 98.93617 | NP_624982.1:c.12G>A | | No | G:C>A:T | SCO0673 | hypothetical protein |
| | 1134186 | SNV | C | T | 156 | 156 | 100 | NP_625368.1:c.30C>T | | No | G:C>A:T | SCO1074 | hypothetical protein |
| | 1467045 | SNV | G | A | 164 | 165 | 99.39394 | NP_625671.1:c.717C>T | | No | G:C>A:T | SCO1388 | mannose-1-phosphate guanyltransferase |
| | 1503926 | SNV | C | T | 179 | 183 | 97.81421 | NP_625691.1:c.394G>A | NP_625691.1:p.Val132Met | Yes | G:C>A:T | SCO1409 | hypothetical protein |
| | 1615858 | SNV | C | T | 145 | 146 | 99.31507 | NP_733536.1:c.559C>T | NP_733536.1:p.His187Tyr | Yes | G:C>A:T | SCO1511 | hypothetical protein |
| | 1811018 | SNV | G | T | 185 | 185 | 100 | NP_625965.1:c.369C>A | | No | G:C>T:A | SCO1691 | TetR family transcriptional regulator |
| | 1906862 | SNV | G | T | 150 | 150 | 100 | NP_626052.1:c.367C>A | | No | G:C>T:A | ppnK | inorganic polyphosphate/ATP-NAD kinase |
| | 2119778 | SNV | G | A | 140 | 141 | 99.29078 | NP_626243.1:c.19G>A | NP_626243.1:p.Val7Ile | Yes | G:C>A:T | SCO1981 | hypothetical protein |
| | 2458948^245894 | Insertion | - | C | 83 | 138 | 60.14493 | | | - | Insertion | | |
| | 2489834 | SNV | C | G | 159 | 160 | 99.375 | NP_626565.1:c.221G>C | NP_626565.1:p.Gly74Ala | Yes | G:C>C:G | SCO2318 | glycosyl transferase |
| | 2884357 | SNV | G | C | 170 | 172 | 98.83721 | NP_626888.1:c.274G>C | NP_626888.1:p.Ala92Pro | Yes | G:C>C:G | SCO2652 | hypothetical protein |
| | 2968251 | SNV | C | A | 154 | 154 | 100 | NP_626956.1:c.1936G>T | NP_626956.1:p.Glu646* | Yes | G:C>T:A | SCO2723 | ABC transporter ATP-binding protein |
| | 3345283 | SNV | C | A | 174 | 178 | 97.75281 | NP_627273.1:c.864C>A | | No | G:C>T:A | SCO3052 | UDP-glucose 6-dehydrogenase |
| | 4040725 | SNV | G | T | 147 | 147 | 100 | NP_733613.1:c.2323C>A | | No | G:C>T:A | SCO3661 | ATP-dependent protease ATP-binding subunit |
| | 4127767 | SNV | G | T | 51 | 100 | 51 | NP_733616.1:c.212C>A | NP_733616.1:p.Ala71Asp | Yes | G:C>T:A | SCO3754 | ABC transporter |
| | 4148505 | SNV | T | C | 24 | 25 | 96 | | | - | A:T>G:C | | |
| | 4532381 | SNV | T | C | 116 | 118 | 98.30508 | | | - | A:T>G:C | | |
| | 4732128 | SNV | C | G | 171 | 172 | 99.4186 | NP_628492.1:c.2546G>C | NP_628492.1:p.Arg849Pro | Yes | G:C>C:G | SCO4321 | hypothetical protein |
| | 5777386 | SNV | C | T | 164 | 164 | 100 | NP_629446.1:c.489G>A | | No | G:C>A:T | SCO5304 | sensor-like histidine kinase |
| | 5840932 | SNV | C | G | 175 | 183 | 95.62842 | NP_629510.1:c.762C>G | | No | G:C>C:G | SCO5371 | F0F1 ATP synthase subunit alpha |
| | 5960922 | SNV | C | A | 145 | 145 | 100 | | | - | G:C>T:A | | |
| | 6203952 | SNV | C | G | 144 | 144 | 100 | NP_629822.1:c.31C>G | NP_629822.1:p.Pro11Ala | Yes | G:C>C:G | SCO5694 | 1-deoxy-D-xylulose 5-phosphate reductoisomerase |
| | 6371032 | SNV | C | A | 186 | 186 | 100 | NP_629945.1:c.27C>A | | No | G:C>T:A | SCO5822 | DNA topoisomerase IV subunit B |
| | 6629123 | SNV | G | C | 153 | 155 | 98.70968 | NP_630149.1:c.509C>G | NP_630149.1:p.Ala170Gly | Yes | G:C>C:G | SCO6038 | hypothetical protein |
| | 6960345 | SNV | G | C | 173 | 174 | 99.42529 | NP_630397.1:c.1466C>G | NP_630397.1:p.Thr489Ser | Yes | G:C>C:G | SCO6300 | hydrolase |
| | 6972333 | SNV | G | C | 145 | 146 | 99.31507 | NP_630409.1:c.343C>G | NP_630409.1:p.Pro115Ala | Yes | G:C>C:G | SCO6312 | transcriptional regulator |
| | 7009564 | Deletion | G | - | 144 | 145 | 99.31034 | NP_630440.1:c.1329delC | NP_630440.1:p.Ala444fs | Yes | Deletion | SCO6348 | hypothetical protein |
| | 7009566 | SNV | C | T | 144 | 145 | 99.31034 | NP_630440.1:c.1327G>A | NP_630440.1:p.Val443Ile | Yes | G:C>A:T | SCO6348 | hypothetical protein |
| | 7101047 | SNV | G | T | 165 | 165 | 100 | NP_630514.1:c.3465C>A | | No | G:C>T:A | SCO6428 | hypothetical protein |
| | 7271648 | SNV | G | A | 148 | 148 | 100 | NP_630648.1:c.546G>A | | No | G:C>A:T | SCO6568 | ABC transporter |
| | 7547616 | SNV | G | C | 168 | 170 | 98.82353 | NP_630861.1:c.394G>C | NP_630861.1:p.Val132Leu | Yes | G:C>C:G | SCO6789 | fatty oxidation protein |
| | 7559869 | SNV | G | A | 156 | 156 | 100 | NP_630871.1:c.240G>A | | No | G:C>A:T | tdh | L-threonine 3-dehydrogenase |
| | 7566905 | SNV | G | C | 162 | 165 | 98.18182 | NP_630878.1:c.443C>G | NP_630878.1:p.Ala148Gly | Yes | G:C>C:G | SCO6806 | phage integrase |
| | 7766466 | SNV | C | G | 297 | 300 | 99 | NP_631060.1:c.210C>G | | No | G:C>C:G | SCO6995 | protease |
| M1 | 7776421 | SNV | G | A | 320 | 328 | 97.56098 | NP_631068.1:c.752C>T | NP_631068.1:p.Ser251Phe | Yes | G:C>A:T | SCO7003 | hypothetical protein |
| | 1088640 | SNV | C | A | 127 | 128 | 99.21875 | NP_625326.1:c.2475G>T | | No | G:C>T:A | SCO1031 | ABC transporter |
| | 1126417 | SNV | G | A | 71 | 136 | 52.20588 | | | - | G:C>A:T | | |
| | 1166168 | SNV | G | C | 104 | 104 | 100 | NP_625402.1:c.253G>C | NP_625402.1:p.Glu85Gln | Yes | G:C>C:G | SCO1109 | oxidoreductase |
| | 1404674 | SNV | C | T | 150 | 150 | 100 | | | - | G:C>A:T | | |
| | 1432260 | SNV | G | A | 122 | 122 | 100 | | | - | G:C>A:T | | |
| | 1690052 | SNV | C | A | 94 | 96 | 97.91667 | NP_733539.1:c.385G>T | NP_733539.1:p.Ala129Ser | Yes | G:C>T:A | argJ | bifunctional ornithineacetyltransferase/N-acetylglutamate synthase |
| | 1719518 | SNV | G | T | 103 | 103 | 100 | | | - | G:C>T:A | | |
| | 1807393 | SNV | C | T | 86 | 92 | 93.47826 | | | - | G:C>A:T | | |
| | 1926842 | SNV | C | T | 160 | 160 | 100 | NP_626068.1:c.673G>A | NP_626068.1:p.Val225Ile | Yes | G:C>A:T | SCO1798 | ABC transporter ATP-binding protein |
| | 2542975 | SNV | G | A | 109 | 109 | 100 | | | - | G:C>A:T | | |
| | 2613098 | SNV | C | A | 180 | 181 | 99.44751 | | | - | G:C>T:A | | |
| | 2758023 | SNV | C | T | 146 | 146 | 100 | NP_626796.1:c.440G>A | NP_626796.1:p.Gly147Glu | Yes | G:C>A:T | SCO2558 | hypothetical protein |
| | 2864030 | SNV | G | A | 119 | 122 | 97.54098 | NP_626871.1:c.2329G>A | NP_626871.1:p.Val777Met | Yes | G:C>A:T | SCO2635 | aminopeptidase |
| | 2908625 | SNV | G | C | 91 | 96 | 94.79167 | NP_626907.1:c.260G>C | NP_626907.1:p.Gly87Ala | Yes | G:C>C:G | SCO2672 | hypothetical protein |
| | 2990314 | SNV | A | T | 115 | 115 | 100 | | | - | A:T>T:A | | |
| | 2993995 | SNV | G | C | 63 | 96 | 65.625 | NP_626978.1:c.1005G>C | | No | G:C>C:G | SCO2747 | bifunctional carbohydrate binding and transportprotein |
| | 3282758 | SNV | G | C | 229 | 233 | 98.28326 | | | - | G:C>C:G | | |
| | 3583178 | SNV | G | A | 177 | 178 | 99.4382 | NP_733597.1:c.6444G>A | | No | G:C>A:T | SCO3232 | CDA peptide synthetase III |
| | 3585292 | SNV | C | T | 148 | 150 | 98.66667 | NP_627446.1:c.471C>T | | No | G:C>A:T | SCO3234 | phosphotransferase |
| | 3796098 | SNV | G | T | 117 | 119 | 98.31933 | NP_733604.1:c.1366G>T | NP_733604.1:p.Ala456Ser | Yes | G:C>T:A | SCO3434 | DNA polymerase I |
| | 4313821 | SNV | T | G | 299 | 299 | 100 | | | - | A:T>C:G | | |
| | 4486909 | SNV | G | A | 173 | 177 | 97.74011 | NP_628273.1:c.3714C>T | | No | G:C>A:T | SCO4092 | ATP-dependent helicase |
| | 4617566 | SNV | A | G | 206 | 222 | 92.79279 | NP_628383.1:c.293T>C | NP_628383.1:p.Val98Ala | Yes | A:T>G:C | SCO4208 | integral membrane transport protein |
| | 4811070 | SNV | C | T | 144 | 230 | 62.6087 | NP_628563.1:c.235C>T | NP_628563.1:p.His79Tyr | Yes | G:C>A:T | SCO4394 | iron repressor |
| | 5532591 | SNV | C | T | 195 | 197 | 98.98477 | NP_629239.1:c.143C>T | NP_629239.1:p.Thr48Ile | Yes | G:C>A:T | SCO5089 | actinorhodin polyketide synthase |
| | 5775556 | SNV | G | A | 202 | 204 | 99.01961 | NP_629444.1:c.423C>T | | No | G:C>A:T | SCO5302 | integral membrane cell-cycle protein |
| | 6037691 | SNV | C | G | 191 | 192 | 99.47917 | NP_629674.1:c.1064G>C | NP_629674.1:p.Arg355Pro | Yes | G:C>C:G | SCO5540 | hypothetical protein |
| | 7129427 | SNV | A | T | 112 | 112 | 100 | | | - | A:T>T:A | | |
| | 7298296 | SNV | C | G | 89 | 89 | 100 | NP_630666.1:c.867C>G | | No | G:C>C:G | SCO6587 | dehydrogenase |
| | 7406743 | SNV | G | C | 83 | 83 | 100 | NP_630741.1:c.909C>G | NP_630741.1:p.Ser303Arg | Yes | G:C>C:G | SCO6666 | hypothetical protein |
| | 7634733 | SNV | C | A | 78 | 78 | 100 | NP_630934.1:c.946G>T | NP_630934.1:p.Gly316Cys | Yes | G:C>T:A | SCO6864 | hypothetical protein |
| | 7645276 | SNV | T | A | 116 | 117 | 99.1453 | NP_630946.1:c.131A>T | NP_630946.1:p.Lys44Met | Yes | A:T>T:A | SCO6876 | hypothetical protein |
| M2 | 7807060 | SNV | G | T | 51 | 52 | 98.07692 | NP_631085.1:c.40G>T | NP_631085.1:p.Gly14Trp | Yes | G:C>T:A | SCO7021 | hypothetical protein |
| | 1072216..1072217 | MNV | GA | TC | 117 | 121 | 96.69421 | NP_625312.1:c.328_329delTCinsGA | NP_625312.1:p.Ser110Asp | Yes | G:C>T:A<br>A:T>C:G | SCO1016 | hypothetical protein |
| | 1252185 | SNV | C | T | 154 | 161 | 95.65217 | NP_625474.1:c.7C>T | NP_625474.1:p.Arg3Cys | Yes | G:C>A:T | SCO1184 | hypothetical protein |
| | 1344799 | SNV | A | C | 27 | 53 | 50.9434 | NP_625561.1:c.653A>C | NP_625561.1:p.His218Pro | Yes | A:T>C:G | SCO1274 | hypothetical protein |
| | 1592949 | SNV | A | C | 58 | 92 | 63.04348 | | | - | A:T>C:G | | |
| | 1719817 | SNV | T | G | 64 | 120 | 53.33333 | | | - | A:T>C:G | | |
| | 1765203 | SNV | A | G | 38 | 74 | 51.35135 | | | - | A:T>G:C | | |
| | 1878761 | SNV | T | G | 63 | 119 | 52.94118 | | | - | A:T>C:G | | |
| | 1878763 | SNV | A | G | 68 | 117 | 58.11966 | | | - | A:T>G:C | | |
| | 1878792 | SNV | T | C | 62 | 121 | 51.23967 | | | - | A:T>G:C | | |
| | 2112369 | SNV | C | T | 131 | 131 | 100 | NP_626236.1:c.742G>A | NP_626236.1:p.Gly248Ser | Yes | G:C>A:T | SCO1973 | hypothetical protein |
| | 2686414 | SNV | G | T | 179 | 180 | 99.44444 | NP_626736.1:c.1220C>A | NP_626736.1:p.Pro407Gln | Yes | G:C>T:A | SCO2494 | pyruvate phosphate dikinase |
| | 2947470 | SNV | A | T | 281 | 282 | 99.64539 | | | - | A:T>T:A | | |
| | 4148246 | SNV | A | C | 11 | 21 | 52.38095 | | | - | A:T>C:G | | |
| | 4148252 | SNV | A | C | 10 | 20 | 50 | | | - | A:T>C:G | | |
| | 4377204^437720 | Insertion | - | G | 137 | 174 | 78.73563 | | | - | Insertion | | |
| | 4430625 | SNV | A | C | 65 | 130 | 50 | | | - | A:T>C:G | | |
| | 4663287 | SNV | G | T | 158 | 174 | 90.8046 | NP_628428.1:c.39G>T | | No | G:C>T:A | SCO4254 | hypothetical protein |
| | 4770844 | SNV | A | G | 240 | 240 | 100 | NP_628526.1:c.296T>C | NP_628526.1:p.Leu99Pro | Yes | A:T>G:C | SCO4356 | hypothetical protein |
| | 4880312 | SNV | C | T | 276 | 281 | 98.22064 | | | - | G:C>A:T | | |
| | 5255390 | SNV | A | C | 103 | 191 | 53.9267 | | | - | A:T>C:G | | |
| | 5612258 | SNV | A | G | 36 | 70 | 51.42857 | | | - | A:T>G:C | | |
| | 5612260 | SNV | T | G | 37 | 70 | 52.85714 | | | - | A:T>C:G | | |
| | 5612263 | SNV | C | G | 47 | 91 | 51.64835 | | | - | G:C>C:G | | |
| | 5830280 | SNV | T | C | 129 | 256 | 50.39063 | | | - | A:T>G:C | | |
| | 5871569 | SNV | C | T | 225 | 226 | 99.55752 | NP_629538.1:c.957C>T | | No | G:C>A:T | SCO5399 | acetyl-CoA acetyltransferase |
| | 6105201 | SNV | C | A | 169 | 169 | 100 | NP_629737.1:c.334C>A | NP_629737.1:p.Leu112Ile | Yes | G:C>T:A | SCO5603 | hypothetical protein |
| | 6113723 | SNV | G | C | 192 | 193 | 99.48187 | NP_629746.1:c.2170G>C | NP_629746.1:p.Val724Leu | Yes | G:C>C:G | SCO5612 | ATP binding protein |
| | 6598610 | SNV | A | C | 50 | 99 | 50.50505 | | | - | A:T>C:G | | |
| | 6935489 | SNV | C | T | 142 | 145 | 97.93103 | NP_630376.1:c.983C>T | NP_630376.1:p.Pro328Leu | Yes | G:C>A:T | SCO6278 | integral membrane transport protein |
| | 7633600 | SNV | C | G | 193 | 193 | 100 | NP_630933.1:c.220G>C | NP_630933.1:p.Ala74Pro | Yes | G:C>C:G | SCO6863 | hypothetical protein |
| M3 | 7777209 | SNV | G | C | 153 | 154 | 99.35065 | | | - | G:C>C:G | | |
| | 527567 | SNV | G | A | 160 | 162 | 98.76543 | NP_624811.1:c.272C>T | NP_624811.1:p.Pro91Leu | Yes | G:C>A:T | SCO0494 | iron-siderophore binding lipoprotein |
| | 533965 | SNV | C | T | 114 | 114 | 100 | NP_624818.1:c.1800G>A | | No | G:C>A:T | SCO0501 | hypothetical protein |
| | 738352^738353 | Insertion | - | G | 71 | 106 | 66.98113 | | | - | Insertion | | |
| | 5056054 | SNV | C | G | 188 | 189 | 99.4709 | NP_628793.1:c.227C>G | NP_628793.1:p.Thr76Ser | Yes | G:C>C:G | SCO4632 | ATP/GTP binding protein |
| | 5257242 | SNV | G | A | 172 | 173 | 99.42197 | | | - | G:C>A:T | | |
| | 6370279^637028 | Insertion | - | G | 48 | 75 | 64 | | | - | Insertion | | |
| M4 | 7595845 | SNV | G | A | 113 | 150 | 75.33333 | NP_630898.1:c.2711C>T | NP_630898.1:p.Ala904Val | Yes | G:C>A:T | SCO6827 | polyketide synthase |
| | 2462887 | Deletion | G | - | 67 | 108 | 62.03704 | | | - | Deletion | rnpB | |
| | 2961862 | SNV | G | T | 126 | 126 | 100 | NP_626951.1:c.153C>A | | No | G:C>T:A | SCO2718 | hypothetical protein |
| | 5203768 | SNV | T | C | 143 | 147 | 97.27891 | NP_628940.1:c.29A>G | NP_628940.1:p.Asp10Gly | Yes | A:T>G:C | SCO4782 | hypothetical protein |
| M5 | 6769937 | SNV | G | C | 33 | 59 | 55.9322 | NP_630271.1:c.530G>C | NP_630271.1:p.Arg177Pro | Yes | G:C>C:G | SCO6167 | proline rich protein membrane protein |
| | 454451^454452 | Insertion | - | C | 192 | 274 | 70.07299 | | | - | Insertion | | |
| | 489245 | Deletion | G | - | 206 | 208 | 99.03846 | NP_624786.1:c.632delG | NP_624786.1:p.Arg211fs | Yes | Deletion | SCO0467 | hypothetical protein |
| | 606363 | SNV | G | T | 168 | 174 | 96.55172 | NP_624876.1:c.193C>A | NP_624876.1:p.Arg65Ser | Yes | G:C>T:A | SCO0563 | hypothetical protein |
| | 681473 | SNV | C | G | 152 | 155 | 98.06452 | NP_624950.1:c.391G>C | NP_624950.1:p.Glu131Gln | Yes | G:C>C:G | SCO0640 | hypothetical protein |
| | 1251111 | SNV | G | C | 126 | 132 | 95.45455 | NP_625473.1:c.108G>C | | No | G:C>C:G | SCO1183 | hypothetical protein |
| | 1562784 | SNV | C | G | 131 | 132 | 99.24242 | NP_625746.1:c.737G>C | NP_625746.1:p.Cys246Ser | Yes | G:C>C:G | SCO1465 | hypothetical protein |
| | 1670045 | SNV | G | T | 136 | 137 | 99.27007 | NP_625836.1:c.628C>A | NP_625836.1:p.Leu210Met | Yes | G:C>T:A | SCO1558 | ABC transporter permease |
| | 1765203^176520 | Insertion | - | G | 54 | 89 | 60.67416 | | | - | Insertion | | |
| | 1984265 | SNV | G | A | 155 | 156 | 99.35897 | NP_626119.1:c.244G>A | NP_626119.1:p.Gly82Arg | Yes | G:C>A:T | SCO1851 | cob(I)yrinic acid a,c-diamideadenosyltransferase |
| | 1988841 | SNV | A | C | 103 | 105 | 98.09524 | NP_626123.1:c.373A>C | NP_626123.1:p.Thr125Pro | Yes | A:T>C:G | SCO1856 | precorrin-6Y C5,15-methyltransferase |
| | 2295007 | SNV | G | A | 172 | 172 | 100 | NP_626389.1:c.1126G>A | NP_626389.1:p.Gly376Ser | Yes | G:C>A:T | SCO2133 | hypothetical protein |
| | 2646572 | SNV | A | G | 181 | 182 | 99.45055 | NP_626703.1:c.79T>C | | No | A:T>G:C | SCO2460 | hypothetical protein |
| | 3008164 | SNV | T | A | 89 | 89 | 100 | | | - | A:T>T:A | | |
| | 4377204^437720 | Insertion | - | G | 106 | 156 | 67.94872 | | | - | Insertion | | |
| | 4564696 | SNV | C | G | 187 | 189 | 98.9418 | NP_628325.1:c.117C>G | | No | G:C>C:G | SCO4148 | ABC transporter ATP-binding protein |
| | 4658248 | SNV | G | T | 130 | 130 | 100 | NP_628424.1:c.1348G>T | NP_628424.1:p.Gly450Trp | Yes | G:C>T:A | SCO4250 | hypothetical protein |
| | 4880904 | SNV | C | T | 129 | 138 | 93.47826 | NP_733638.1:c.71G>A | NP_733638.1:p.Gly24Asp | Yes | G:C>A:T | SCO4458 | lipoprotein |
| | 4940282 | SNV | A | C | 200 | 202 | 99.0099 | NP_628682.1:c.580T>G | NP_628682.1:p.Trp194Gly | Yes | A:T>C:G | SCO4518 | hypothetical protein |
| | 6503843 | SNV | G | T | 131 | 132 | 99.24242 | NP_630050.1:c.80C>A | NP_630050.1:p.Pro27His | Yes | G:C>T:A | SCO5933 | hypothetical protein |
| | 6597353 | SNV | A | T | 193 | 196 | 98.46939 | NP_630127.1:c.312A>T | NP_630127.1:p.Glu104Asp | Yes | A:T>T:A | SCO6014 | cationic amino acid transporter |
| | 6826638 | SNV | C | T | 174 | 177 | 98.30508 | | | - | G:C>A:T | | |
| | 6994242 | SNV | G | C | 155 | 161 | 96.27329 | NP_630426.1:c.192C>G | | No | G:C>C:G | SCO6334 | transcriptional regulator |
| | 7163598 | SNV | G | A | 189 | 189 | 100 | | | - | G:C>A:T | | |
| | 7198042 | SNV | C | A | 152 | 153 | 99.34641 | NP_630588.1:c.111C>A | NP_630588.1:p.Asp37Glu | Yes | G:C>T:A | SCO6506 | gas vesicle protein |
| | 7237535 | SNV | G | T | 127 | 127 | 100 | NP_630625.1:c.322G>T | NP_630625.1:p.Glu108* | Yes | G:C>T:A | SCO6544 | hypothetical protein |

| | | | | | | | | | | | | | |
|---|---|---|---|---|---|---|---|---|---|---|---|---|---|
| | 7671689 | SNV | G | A | 176 | 176 | 100 | NP_630976.1:c.355G>A | NP_630976.1:p.Val119Met | Yes | G:C>A:T | SCO6907 | DNA ligase |
| | 7755660 | SNV | C | G | 96 | 101 | 95.0495 | NP_631053.1:c.1125G>C | | No | G:C>C:G | SCO6988 | oxidoreductase |
| M6 | 7783012 | Deletion | G | - | 73 | 73 | 100 | NP_631073.1:c.352delG | NP_631073.1:p.Val118fs | Yes | Deletion | SCO7008 | ABC transporter ATP-binding protein |
| | 466074 | SNV | A | C | 53 | 55 | 96.36364 | NP_624768.1:c.85A>C | NP_624768.1:p.Ile29Leu | Yes | A:T>C:G | SCO0447 | MarR family regulatory protein |
| | 515777 | SNV | C | G | 118 | 119 | 99.15966 | NP_624809.1:c.9144G>C | | No | G:C>C:G | SCO0492 | peptide synthetase |
| | 1467501 | SNV | T | A | 146 | 148 | 98.64865 | NP_625671.1:c.261A>T | | No | A:T>T:A | SCO1388 | mannose-1-phosphate guanyltransferase |
| | 1765203^176520 | Insertion | - | G | 51 | 84 | 60.71429 | | | - | Insertion | | |
| | 1792889 | SNV | G | A | 117 | 117 | 100 | NP_625946.1:c.2045G>A | NP_625946.1:p.Arg682Gln | Yes | G:C>A:T | SCO1671 | hypothetical protein |
| | 2077625 | SNV | G | A | 171 | 173 | 98.84393 | | | - | G:C>A:T | | |
| | 2139959 | SNV | G | C | 101 | 101 | 100 | NP_626262.1:c.765C>G | NP_626262.1:p.Tyr255* | Yes | G:C>C:G | SCO2001 | hypothetical protein |
| | 3137890 | SNV | G | A | 197 | 200 | 98.5 | NP_627111.1:c.230G>A | NP_627111.1:p.Arg77Gln | Yes | G:C>A:T | SCO2883 | cytochrome P450 |
| | 3155674 | SNV | G | A | 192 | 193 | 99.48187 | NP_627128.1:c.219C>T | | No | G:C>A:T | SCO2902 | deoxyribonucleotide triphosphatepyrophosphatase |
| | 3277441 | SNV | G | C | 198 | 199 | 99.49749 | NP_627227.1:c.2280C>G | NP_627227.1:p.Asp760Glu | Yes | G:C>C:G | SCO3005 | preprotein translocase subunit SecA |
| | 3560736 | SNV | G | C | 38 | 75 | 50.66667 | NP_627443.1:c.17402G>C | NP_627443.1:p.Arg5801Pro | Yes | G:C>C:G | SCO3230 | CDA peptide synthetase I |
| | 3910675 | SNV | G | A | 184 | 184 | 100 | NP_627739.1:c.783C>T | | No | G:C>A:T | SCO3541 | DNA polymerase III subunit delta' |
| | 4377204^437720 | Insertion | - | G | 86 | 124 | 69.35484 | | | - | Insertion | | |
| | 4700228 | SNV | C | A | 197 | 198 | 99.49495 | NP_628456.1:c.316G>T | NP_628456.1:p.Gly106Trp | Yes | G:C>T:A | SCO4284 | N-acetylglucosamine-6-phosphate deacetylase |
| | 5611647 | SNV | C | T | 131 | 131 | 100 | NP_629313.1:c.543G>A | | No | G:C>A:T | SCO5165 | hydrolase |
| | 6052711 | SNV | C | T | 140 | 143 | 97.9021 | NP_629687.1:c.699C>T | | No | G:C>A:T | SCO5553 | 3-isopropylmalate dehydratase large subunit |
| | 6613228 | SNV | C | G | 157 | 160 | 98.125 | NP_630137.1:c.2073G>C | | No | G:C>C:G | SCO6026 | fatty acid oxidation complex alpha-subunit |
| | 7224671 | SNV | C | G | 104 | 106 | 98.11321 | NP_630614.1:c.283G>C | NP_630614.1:p.Gly95Arg | Yes | G:C>C:G | SCO6533 | nitrate reductase subunit delta NarJ |
| W3 | 7799431 | SNV | A | T | 176 | 178 | 98.8764 | | | - | A:T>T:A | | |
